# Dysfunction of Torr causes a Harlequin-type ichthyosis-like phenotype in *Drosophila melanogaster*

**DOI:** 10.1101/734400

**Authors:** Y Wang, M Norum, K Oehl, Y Yang, R Zuber, J Yang, JP Farine, N Gehring, M Flötenmeyer, J.-F Ferveur, B Moussian

## Abstract

Prevention of desiccation is a constant challenge for terrestrial organisms. Land insects have an extracellular coat, the cuticle, that plays a major role in protection against exaggerated water loss. Here, we report that the ABC transporter Torr - a human ABCA12 paralog - contributes to the waterproof barrier function of the cuticle in the fruit fly *Drosophila melanogaster*. We show that the reduction or elimination of Torr function provokes rapid desiccation. Torr is also involved in defining the inward barrier against xenobiotics penetration. Consistently, the amounts of cuticular hydrocarbons that are involved in cuticle impermeability decrease markedly when Torr activity is reduced. GFP-tagged Torr localises to membrane nano-protrusions within the cuticle, likely pore canals. This suggests that Torr is mediating the transport of cuticular hydrocarbons (CHC) through the pore canals to the cuticle surface. The envelope, which is the outermost cuticle layer constituting the main barrier, is unaffected in *torr* mutant larvae. This contrasts with the function of Snu, another ABC transporter needed for the construction of the cuticular inward and outward barriers, that nevertheless is implicated in CHC deposition. Hence, Torr and Snu have overlapping and independent roles to establish cuticular resistance against transpiration and xenobiotic penetration. The *torr* deficient phenotype parallels the phenotype of Harlequin ichthyosis caused by mutations in the human *abca12* gene. Thus, it seems that the cellular and molecular mechanisms of lipid barrier assembly in the skin are conserved in vertebrates in invertebrates.

**Author Summary:** As in humans, lipids on the surface of the skin of insects protect the organism against excessive water loss and penetration of potentially harmful substances. During evolution, a greasy surface was indeed an essential trait for adaptation to life outside a watery environment. Here, we show that the membrane-gate transporter Torr is needed for the deposition of barrier lipids on the skin surface in the fruit fly *Drosophila melanogaster* through extracellular nano-tubes, called pore canals. In principle, the involvement of Torr parallels the scenario in humans, where the membrane-gate transporter ABCA12 is implicated in the construction of the lipid-based stratum corneum of the skin. In both cases, mutations in the genes coding for the respective transporter cause rapid water-loss and are lethal soon after birth. We conclude that the interaction between the organism and the environment obviously implies an analogous mechanism of barrier formation and function in vertebrates and invertebrates.

## Introduction

To avoid desiccation, animals build a lipid-based barrier on their outer surface. In mammals, the stratum corneum (SC), which is the uppermost skin layer, consists of ceramides that are either free or bound to so-called cornified envelope (CE) proteins (Rabionet, Gorgas et al., 2014). Failure to form the ceramide matrix by mutations in the genes coding for ceramide-related enzymes causes transepidermal water loss in mice and humans. Insects are covered by a stratified extracellular matrix (ECM) consisting of the innermost chitinous procuticle, the upper protein-lipid epicuticle and the outermost envelope produced by the underlying epidermis (Moussian, 2010). The envelope constitutes the first water- and xenobiotics repellent barrier and is mainly composed of free and bound lipids (Blomquist & Bagneres, 2010). Biosynthesis of these lipid compounds involves enzymes acting especially in oenocytes that are tightly associated with epidermal cells (Ferveur, 1997, Gutierrez, Wiggins et al., 2007). In the fruit fly *Drosophila melanogaster*, CHC production in oenocytes, indeed, is sufficient to protect the animal against desiccation (Wicker-Thomas, Garrido et al., 2015). While some of the molecules and responsible genes involved have been identified (Qiu, Tittiger et al., 2012), the molecular mechanisms of the lipid-based barrier organisation are not well understood.

Recently, we identified and characterised the function of the ABC transporter Snustorr (Snu) in constructing an inward and outward barrier in the *D. melanogaster* cuticle (Zuber, Norum et al., 2018). Snu is needed for correct localisation of the extracellular protein Snustorr-snarlik (Snsl) at the tips of the pore canals that serve as routes for lipid transport through the procuticle. In *snu* mutant larvae, the envelope is incomplete causing rapid water loss and larval death. Moreover, the cuticle of these animals is permeable to xenobiotics indicating that Snu activity is required for the construction of both the outward and inward barrier. The function of Snu is evolutionary conserved given that its homolog LmABCH-9C in the migratory locust *Locusta migratoria* is necessary to build the cuticular desiccation barrier (Yu, Wang et al., 2017). Consistently, in the red flour beetle *Tribolium castaneum*, the putative Snu orthologue TcABCH-9C has been reported to control the amounts of cuticle lipids (Broehan, Kroeger et al., 2013).

Snu and its putative orthologues are half-type ABC transporters that need dimerisation with other half-type transporters in order to be active: they either form homo- or heterodimers. The genome of many insects also harbours two other genes coding for H-type ABC transporters, ABCH-9A and ABCH-9B (Bretschneider, Heckel et al., 2016, Broehan et al., 2013, Liu, Zhou et al., 2011, Pignatelli, Ingham et al., 2018, Qi, Ma et al., 2016, Yu et al., 2017). While insects have three ABCH transporters, other arthropods such as crustaceans and arachnids have multiple copies of this class of transporters (Dermauw, Osborne et al., 2013). According to phylogenetic analyses, the common ancestor of insects and crustacean had one ABCH type transporter: CG11147/LmABCH-9A, which is likely the primary ancestral protein of this transporter family in insects. The CG11147 coding gene is expressed in the digestive tract during embryogenesis and is therefore unlikely to be required for cuticle formation. The ABCH-9B coding gene, by contrast, is expressed in the embryonic epidermis that produced the larval cuticle. Here, we have focussed our study on the function of ABCH-9B transporters during cuticle differentiation in *D. melanogaster*.

## Results

### CG33970 codes for a ABCH-type transporter related to the human ABCA transporters

During our initial screening procedure, we discovered a candidate gene (*CG33970*) affecting fly resistance to desiccation. Prior to functional characterisation of CG33970, we examined the organisation of the corresponding locus. According to flybase (flybase.org), the *CG33970* locus gives rise to three alternative transcripts (Fig. 1). One transcript yields a short protein, which lacks the 287 N-terminal amino acids of the proteins encoded by the two other transcripts. Since these two transcripts differ only in non-coding regions, they produce identical proteins. The respective protein sequence has 777 residues with an ATPase domain in the N-terminal half of the protein and a transmembrane domain in the C-terminal half of the protein. Thus, CG33970 belongs to the group of half-type ATP-binding cassette (ABC) transporters. Using specific primers, we confirmed the expression of the predicted long and the short transcripts by qPCR.

**Figure 1.**
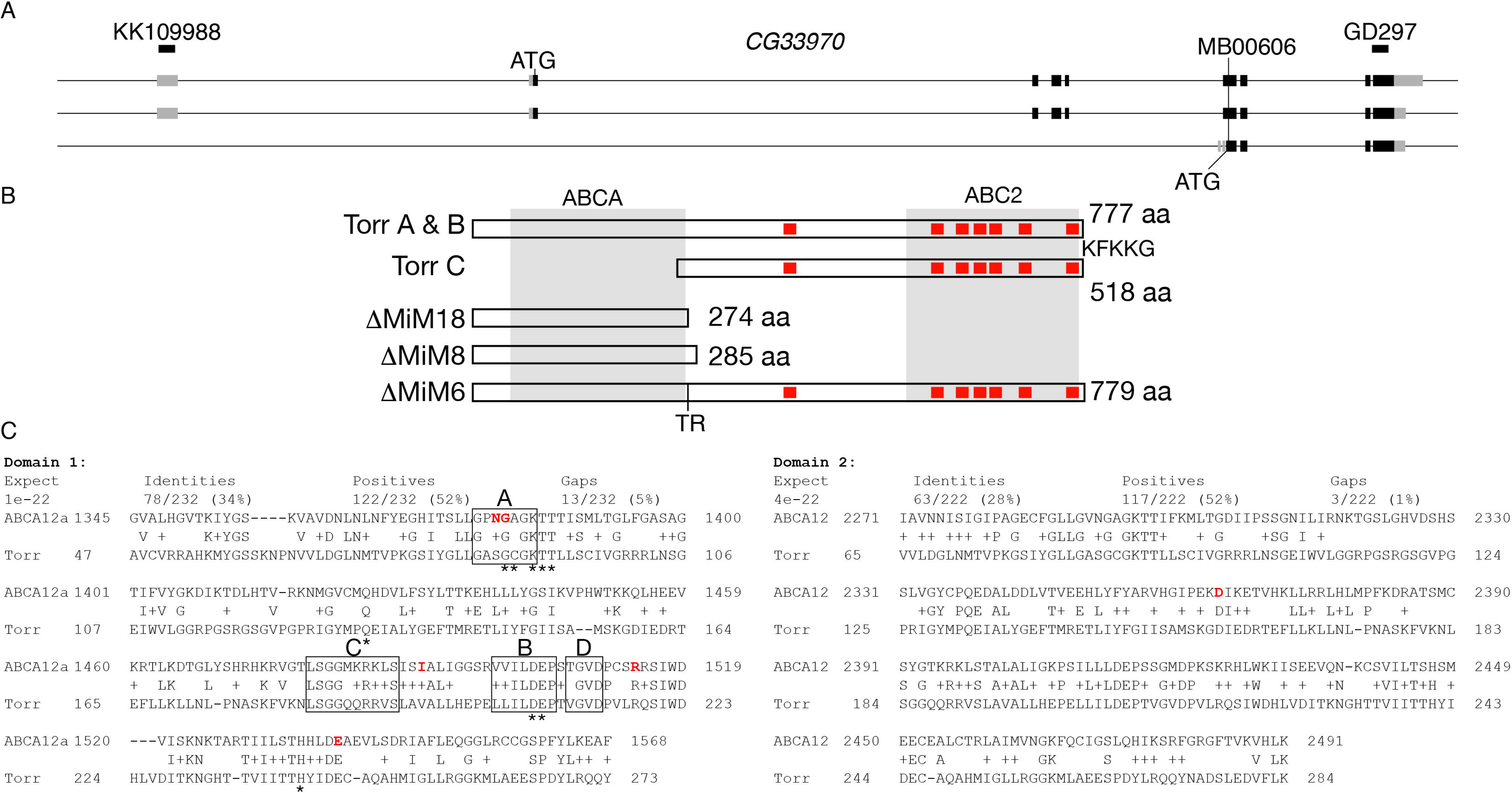
Torr is a half-type ABC transporter. The *CG33970/torr* locus yields three types of transcripts (A), two long and one short mRNAs. The long transcripts are both translated to a 777 residues containing protein with an ABCH domain in its N-terminal half, and an ABC2 domain and six transmembrane domains in its C-terminal half (B). Based on this composition, Torr can be assigned to the class of half-type ABC transporters. In addition, the C-terminus has an ER retention signal (KFKKG). The transposon MB00606 in the 8^th^ exon was excised to obtain alleles coding for dysfunctional proteins. Three lethal excision alleles were sequenced, ΔMiM18, ΔMiM8 and ΔMiM6. These three alleles have insertions of 4 (ΔMiM18), 8 (ΔMiM8) or 6 bp (ΔMiM6) at the position of transposon insertion (in codon number 273). The ΔMiM18 and ΔMiM8 insertions cause frame-shifts resulting in stop codons soon downstream of the insertion site in turn yielding proteins that are devoid of the ABC2 domain. Through the ΔMiM6 insertion, the protein gains two amino acids (TR) in the region linking the ABCH and the ABC2 domains. Alignment of the full-length Torr protein with human ABCA3 and ABCA12 (C). The sequence of the full length Torr protein was blasted against human proteins in the NCBI database using the BlastP software (https://blast.ncbi.nlm.nih.gov/). The Torr ABC cassette domain displays significant homology to both of the respective domains (domain 1 and 2) in all human ABCA transporters. Here, we show the alignment of these domains in ABCA3 and ABCA12 (isoform a). Those amino acids mutated on ABCA12 variants are highlighted in red (Akiyama, 2010). The respective amino acids are marked in orange in the ABCA3 sequence. Not all of these residues are conserved in this sequence.

To learn about the potential molecular function of CG33790, using the full-length CG33790 protein, we searched the NCBI database for human homologous sequences that have been functionally characterised. The most homologous sequences were ABCA-type transporters including ABCA3 and ABCA12 (Fig. 1). ABCA-type transporters are full transporters that are mainly involved in lipid transport across membranes. Hence, it is possible that CG33790 is implicated in lipid transport in *D. melanogaster*.

### Knock-down of CG33970/torr causes post-embryonic desiccation

To investigate the function of the ABC transporter CG33970, we suppressed the expression of all transcripts either in the epidermis or ubiquitously by targeting the UAS-driven transcription of the hairpin RNAs (hpRNA) GD297 (Fig. 1) either with *69B*-Gal4 (Fig. 2) or with the *da*/*7063*-Gal4 drivers, respectively. These larvae became slack just after hatching and they died. Larvae ubiquitously expressing the hpRNA KK109988 that is directed against the long transcripts eventually hatched, but dried out and died about 11 minutes after hatching (Table 1). In general, larvae expressing hpRNA against *CG33970* did not show any obvious morphological defect (Fig. 2). Addition of halocarbon oil to newly hatched *CG33970*^*da/7063-IR*^ knockdown larvae rescued their lethality (Table 1). The “drying-out” phenotype prompted us to name *CG33970 torr* (*torr*, Swedish for dry). In summary, lethality and loss of turgidity just after hatching was induced when hpRNAs against *torr* were expressed under the control of the epidermal *69B*-Gal4 or *da*/*7063*-Gal4 drivers.

**Table 1.** Torr function prevents desiccation

| *treatment* | *wild-type* | *torr<sup>ubIR</sup>* |
| --- | --- | --- |
| Survival time on agar plate | >2h (n=13) | 11min +/- 1.7 (n=10) |
| Survival time on glass & oil | >2h (n=11) | >2h (n=11) |

On agar plates, newly hatched wild-type larvae and larvae ubiquitously (using *da/7063-Gal4*) expressing hpRNA (KK109988) against *torr* (*torr<sup>ubIR</sup>*) were observed for at least two hours (>2h). The second experiment was done on glass because the halocarbon oil would spread on an agar substrate exposing the larvae to the air.

**Figure 2.**
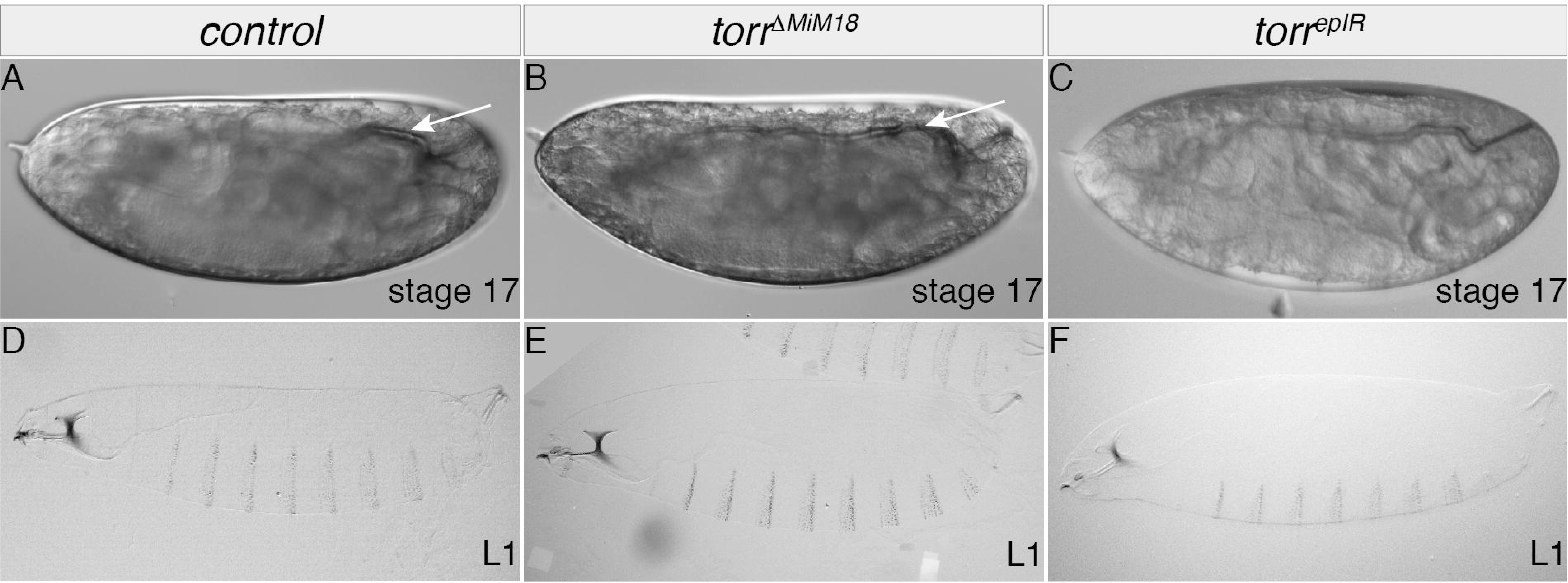
Mutations in the torr gene are lethal. Wild-type stage 17 embryos ready to hatch fill the egg space (A). They have gas-filled tracheae (arrow). Stage 17 embryos homozygous for the *torr* allele *ΔMiM18* (B) or expressing the hpRNA GD297 *torr* in the epidermis using *69B*-Gal4 (C, *torr*^*epIR*^) appear to be normal. Like wild-type 1^st^ instar larvae (D), *torr* mutant 1^st^ instar larvae (E) and 1^st^ instar larvae with reduced *torr* expression (F) hatch, but die soon thereafter (see also Table 1).

Of note, the identical phenotypes caused by expression of KK109988 and GD297 suggests that a possible down-regulation of the potential KK109988 off-target *CG11147* (coding for ABCH-9A, predicted by VDRC) did not contribute to the phenotypes (see discussion).

To confirm the subtle phenotype caused by *torr* reduced expression, we generated stable *torr* mutant alleles by imprecise excision of the *Minos* transposon element *Mi00606* inserted into the exon 7 of the isoforms A and B common coding sequence (see materials & methods and Fig. 1). The line segregating the frame shift mutation *torr*^Δ*MiM18*^ (Fig. 1, additional 4bp, protein length 274 amino acids before a premature stop codon) was used for detailed phenotypic analyses in the following. Larvae homozygous for *torr*^Δ*MiM18*^ did not show any obvious phenotype (Fig. 2). Ubiquitous or epidermal expression of a C-terminally GFP-tagged version of the long isoform of Torr (Torr-GFP) in *torr*^Δ*MiM18*^ larvae using the UAS/Gal4 system (*69B*-Gal4 and *da*/*7063*-Gal4) rescued the lethality and animals survived to adulthood. This result argues that the full-length Torr protein is sufficient for outward barrier construction in the *D. melanogaster* cuticle.

In addition to the embryonic and larval skin, we sought to examine Torr function in a simple cuticle tissue. For this purpose, we suppressed the expression of *torr* in the developing wing using the wing specific *nub*-Gal4 driver to express the *torr*-specific hairpin RNAs. The resulting wings did not show any obvious morphological defect and the respective flies survived and did not desiccate.

### Penetration resistance depends on CG33970 function

The dehydration phenotype of larvae with down-regulated *torr* expression (*torr*^*epIR*^ and *torr*^*ubIR*^) suggests an increased permeability of their cuticle. To test this hypothesis, we incubated these larvae in Eosin Y in a penetration assay (Wang, Carballo et al., 2017, Wang, Yu et al., 2016) (Fig. 3). In these larvae, the reduced Torr function did not cause an abnormal Eosin Y uptake at room temperature (22°C), whereas in *snu* mutant larvae Eosin Y penetrates the tissue in the same assay. At 40°C, however, the cuticle of larvae with reduced *torr* expression was permeable to Eosin Y, while it was not in the wild-type cuticle tested under the same condition. Impermeability to Eosin Y was restored in *torr*^Δ*MiM18*^ larvae that ubiquitously express Torr-GFP (Fig. S2). This result indicates that the full-length Torr isoform is responsible for inward barrier function in *D. melanogaster*.

**Figure 3.**
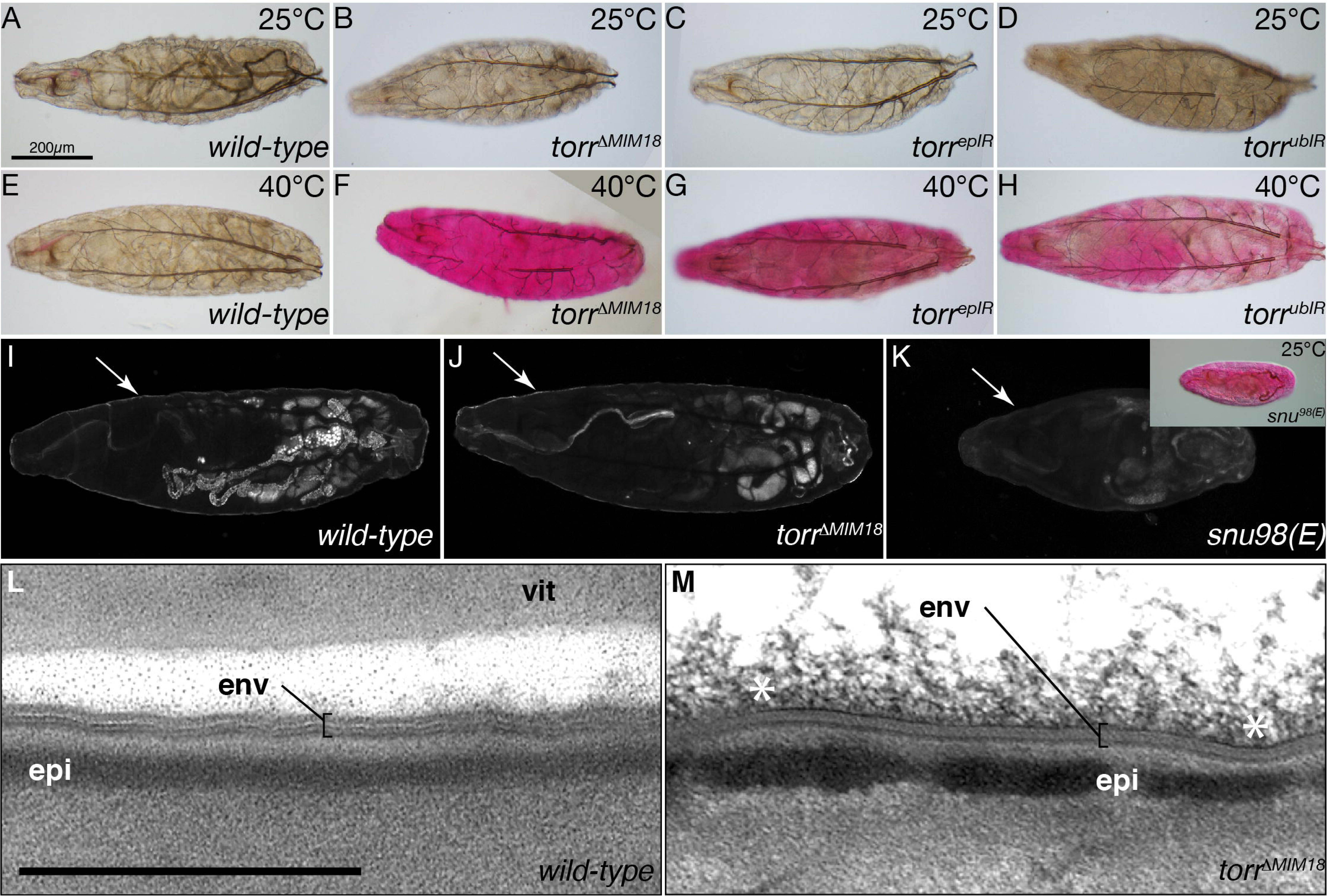
Torr function prevents penetration of xenobiotics. In dye penetration assays, the 1^st^ instar wild-type (A) and *torr* mutant (*torr*^Δ*MiM18*^, B) larvae do not take up Eosin Y at 25°C Down-regulation of *torr* transcripts by epidermal RNAi (*torr*^*epIR*^, *69B*-Gal4 × UAS-KK109988, C) or ubiquitous (*torr*^*ubIR*^, *da/7063*-Gal4 × UAS-KK109988, D) RNAi does not cause dye penetration at 25°C in 1^st^ instar larvae. At 40°C, the cuticle of wild-type 1^st^ instar larvae is impermeable to Eosin Y (D), while in *torr* mutant 1^st^ instar larvae the dye penetrates the cuticle (E). At 25°C, in 1^st^ instar larvae with reduced *torr* transcript levels in the whole body (F) or in the epidermis (G), Eosin Y does not penetrate the cuticle. The cuticle of 1^st^ instar larvae with reduced *torr* transcript levels in the whole body (H) or in the epidermis (I) fails to withstand dye penetration at 40°C. The envelope (arrow) of the wild-type 1^st^ instar larva auto-fluoresces upon excitation with UV light (I). Envelope auto-fluorescence is unchanged in *torr*^Δ*MiM18*^ 1^st^ instar larvae (J). By contrast, envelope auto-fluorescence is reduced in *snu* mutant 1^st^ instar larvae (K). This defect allows Eosin Y penetration into these larvae at 25°C (inset). The envelope (env) of wild-type stage 17 ready-to-hatch embryos consists of alternating electron-lucid and electron-dense sheets in electron-micrographs (L). The envelope of stage 17 ready-to-hatch *torr*^Δ*MiM18*^embryos is normal in electron-micrographs (M). The asterisks (*) mark unspecific material at the surface of the cuticle. vit vitelline membrane. Scale bar 500nm.

We also stained wings with Eosin Y to test whether RNAi-mediated suppression of *torr* expression affected cuticle impermeability in this tissue (Fig. S3). Suppression of *torr* expression under the control of *nub*-Gal4 caused penetration of Eosin Y at a non-permissive temperature. Taken together, these experiments suggest that the suppression of *torr* expression affects both the outward and the inward barrier.

### The envelope of torr mutant larvae is normal

To explore the integrity of the envelope in larvae with reduced or eliminated Torr function, we analysed its auto-fluorescence property when excited with UV light by confocal microscopy (Zuber et al., 2018). The auto-fluorescence of the envelope in *torr*^Δ*MiM18*^ mutant larvae was similar to the signal in wild-type larvae, whereas it was markedly reduced in *snu* mutant larvae (Fig. 3 J-L).

We next examined the potential consequences of Torr dysfunction in the cuticle by transmission electron microscopy of ultrathin sections (Fig. 3 M,N). The cuticle architecture of *torr*^Δ*MiM18*^ homozygous larvae appeared to be normal. Especially, the organisation of the envelope that constitutes a main waterproof barrier was unaffected. Therefore, we conclude that the overall structure of the envelope does not require Torr function.

### Torr localises to the apical plasma membrane independently of Snu

In order to deepen our understanding on Torr function, we determined its sub-cellular localisation. We examined by confocal microscopy the distribution of a chimeric Torr protein with N-terminally fused GFP (GFP-Torr) in L3 larvae. GFP-Torr, which is able to restore cuticle impermeability in *torr*^Δ*MiM18*^ mutant larvae (Fig. S2, see above), localises to the apical plasma membrane of epidermal cells (Fig. 4). Besides a faint uniform localisation at the cell surface, we observed bright GFP-Torr dots. These dots co-localise with dots formed by CD8-RFP (Fig. 4), a membrane marker that also labels membrane protrusions within the cuticle, possibly pore canals (Zuber et al., 2018).

**Figure 4.**
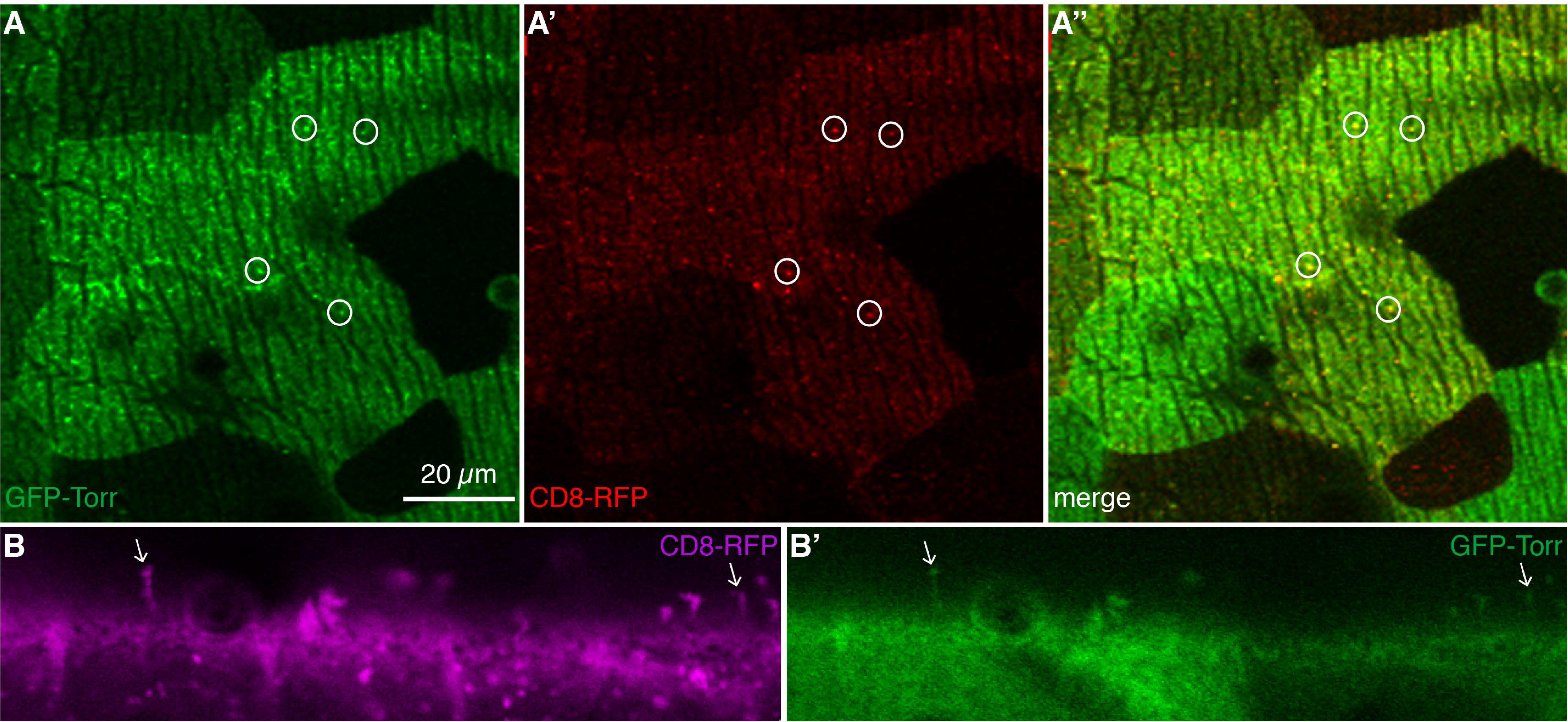
Localization of Snu does not depend on Torr. In stage 16 embryos, Snu-GFP (green) is recruited to the apical plasma membrane of epidermal and gut cells (A,C,C’). This localisation is independent of Torr function (B,D,D’). The apical plasma membrane of gut epithelial cells is marked with Crb (red, C’,D’). A and B live embryos, C and D fixed embryos.

Torr and Snu are both half-type ABC transporters with a C-terminal ER-retention signal (Teasdale & Jackson, 1996). In order to test whether the localisation of Snu in the apical plasma membrane depends on Torr, we observed the distribution of Snu-GFP in *torr*^Δ*MiM18*^ mutant larvae (Fig. 5). In wild-type control larvae and in *torr*^Δ*MiM18*^ mutant larvae, Snu-GFP localized to the apical plasma membrane of epithelial cells. Thus, functional Torr is not required for Snu localisation in the apical plasma membrane. This suggests that Torr and Snu do not interact to form a full ABC transporter.

**Figure 5.**
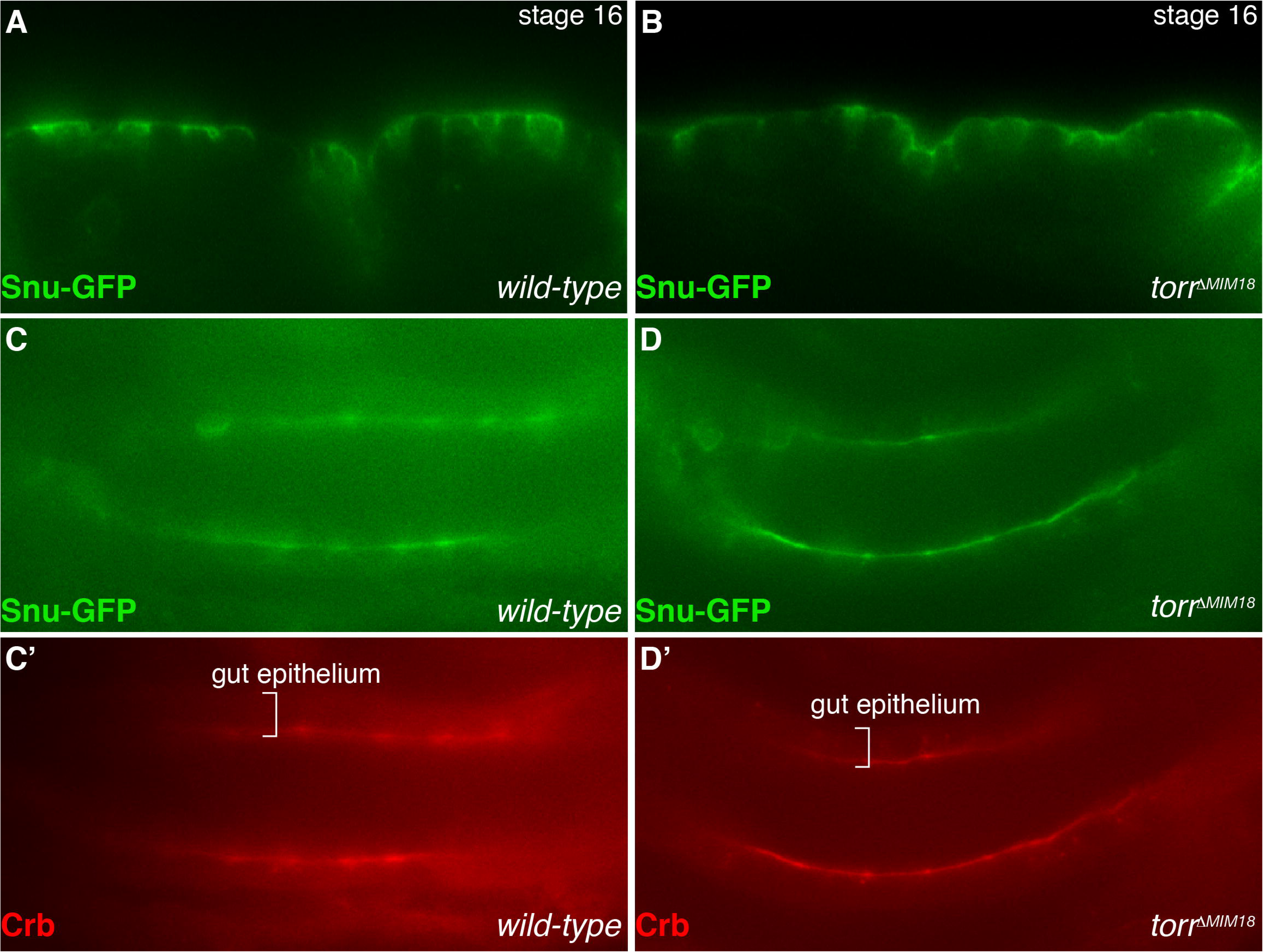
GFP-Torr localizes to pore canals. By confocal microscopy, GFP-Torr (green) is detected at the surface of epidermal cells marked by ridges in the top view (A). Occasionally, the GFP signal accumulates forming dots (circles). These dots co-localises (yellow) with red membrane-marking CD8-RFP dots (A’,A’’). Optical cross-sections confirm that CD8-RFP signals (magenta, B) protruding from the cell surface and marking pore canals coincide (arrows) with GFP-Torr signals (green, B’).

### Torr is required for CHC deposition at the surface of the wing cuticle

CHCs constitute a barrier on the cuticle surface. We quantified and compared their absolute and relative amounts on the wing surface of wild-type flies and flies with reduced *torr* expression after 6 hours post eclosion and at day 5 after eclosion (Fig. 6). When wing-specific RNAi-mediated suppression of *torr* or *snu* were targeted by the nub-Gal4 driver, the resulting wings showed a significant reduction of the total amount of CHCs at both ages compared to the wild type (Dijon 2000) and transgenic control (*nub*-Gal4 × UAS-*CS-2*-IR). Differently, the qualitative variation of the major CHCs types showed no relationship with wing-targeted suppression of *torr* or *snu* expression. Thus, Torr is required for the quantitative deposition of CHCs on the surface of the wing cuticle.

**Figure 6.**
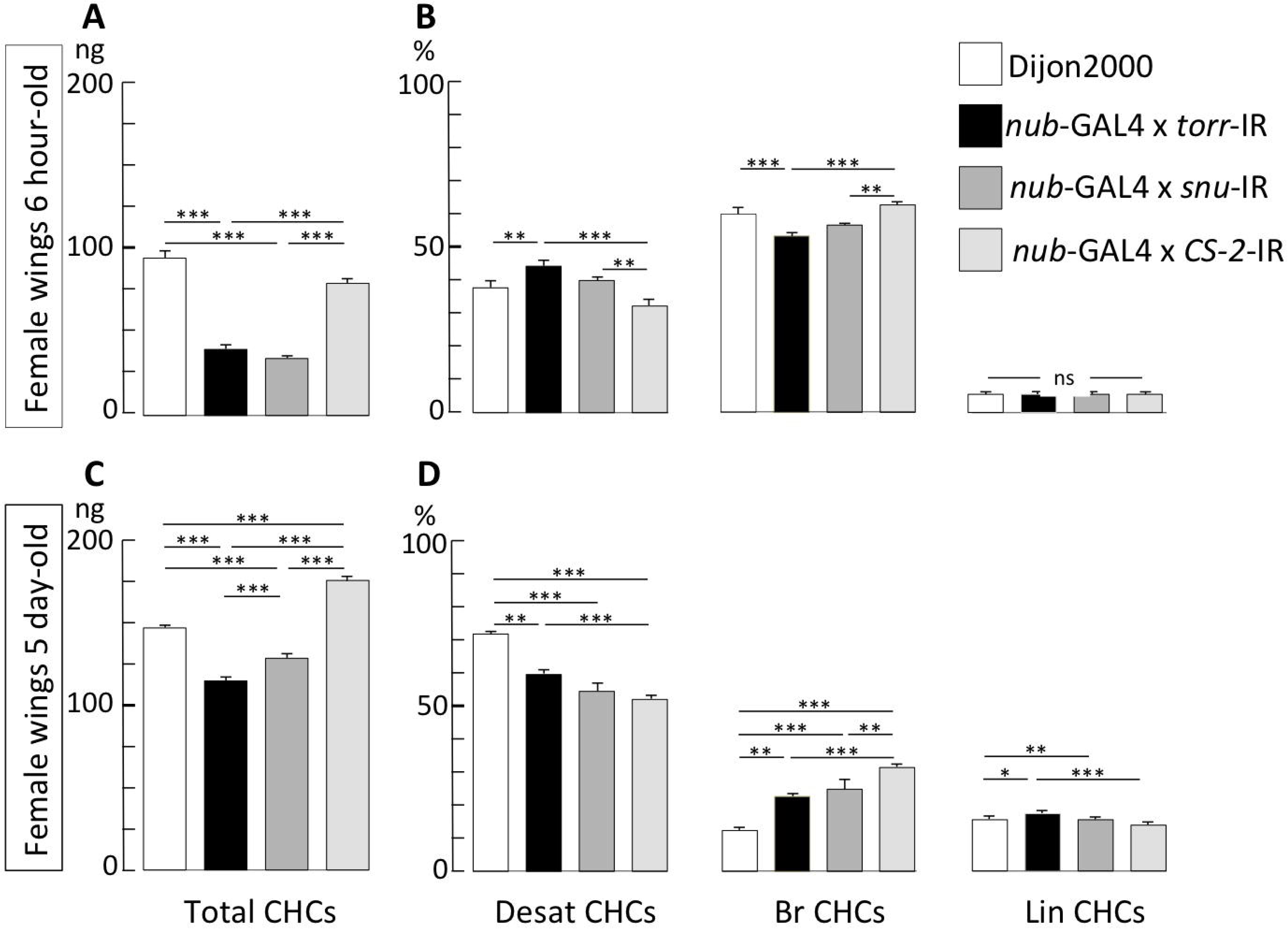
CHC amounts at the wing cuticle surface depend on Torr function. The principal groups of cuticular hydrocarbons (CHCs) measured in pair of wings of individual flies are shown in 6-hour-old (A, B) and 5-day old females (C, D). The genotypes tested are the wild-type Dijon2000 strain (empty bars), the progenies of *nub*-Gal4-RNAi crosses *torr*^*wgIR*^ (KK109988; bars with dark filling), *snu*^*wgIR*^ (bars with medium grey filling), and *CS-2*^*wgIR*^ (bars with light grey filling). The graphs shown on the left (A, C) correspond to the mean (± sem) for the total absolute amount of CHs (Total CHCs) measured in ng. The graphs shown on the right (B, D) correspond to the relative level (%; calculated from the Total CHCs) for the sums of desaturated CHs (Desat CHCs), of branched CHCs (∑Br CHCs), and of linear saturated CHCs (Lin CHCs). Significant differences between genotypes were tested with a Kruskal Wallis test with Conover-Iman multiple pairwise comparisons −α=0.05, with Bonferroni correction; the level of significance between genotypes is indicated as: ***: p<0.001; **: p<0.01; *: p<0.05; ns: not significant. *N*=5 for all genotypes and ages, except for 6 hour-old *snu* nub-KK107544 females: *N*=3.

## Discussion

ABC transporters play important roles in cell and tissue homeostasis by mediating the exchange of solutes and metabolites across membranes. Here we show that Torr, a half-type ABC transporter, is needed for transpiration as well as penetration control in *D. melanogaster*. Moreover, our data indicate that this transporter is directly or indirectly involved in the externalization of CHCs on the cuticular envelope of flies.

### torr codes for a full and a truncated half-type ABC transporter version

The *torr* locus is predicted to have two transcription start sites resulting in three transcripts. Two alternative long mRNA transcribed from the same transcription start site encode the same protein, while a short mRNA starting with exon 7 of the locus encodes a truncated protein lacking the N-terminal NBD. By qPCR, we confirmed the presence of the long and a short *torr* transcripts in the developing embryo and the wing. No truncated version of the other two ABCH-type of transporters, Snu and CG11147, has been predicted or reported. In the literature, however, there are a few reports on truncated ABC transporters in vertebrates. Besides the full length protein, for instance, there are three truncated forms of the human ABCA9 that lack either the C-terminal NBD or even 80% of the full-length protein (Piehler, Kaminski et al., 2002). Two alternative transcription start sites in the human *MRP9* locus lead to mRNAs that code for a full length transporter or a truncated protein that harbours only the NBD, respectively (Bera, Iavarone et al., 2002). In both cases, it is yet unknown whether the truncated proteins have any function. Taken together, our rescue experiments with a GFP-tagged full-length Torr version suggest that the putative truncated Torr protein translated from the short transcript is not essential, while the full-length protein version is needed for viability.

### Insect ABCH transporters act in barrier construction or function

Insects have three ABCH-type transporters, ABCH-9A, −9B and −9C (Broehan et al., 2013, Yu et al., 2017). In *D. melanogaster, T. castaneum* and *L. migratoria* ABCH-9C has been reported to be involved in the construction of the cuticle waterproof barrier (Broehan et al., 2013, Yu et al., 2017, Zuber et al., 2018). In *D. melanogaster* and *L. migratoria*, it was shown that ABCH-9C is also needed to prevent xenobiotics penetration. In this work, we show that Torr, which represents the insect ABCH-9B in *D. melanogaster*, is required to prevent xenobiotics penetration into larvae and wings as well as desiccation in larvae. Thus, two ABCH-type transporters are implicated in the function of the cuticle outward and inward barrier in *D. melanogaster*. The third ABCH transporter CG11147 is expressed in the digestive system and is obviously not important for cuticle barrier function.

### Torr and Snu act in parallel in barrier construction or function

The phenotype of *torr* mutant larvae is, compared to those provoked by mutations in *snu*, rather weak. The envelope structure of *snu* mutant larvae is reduced (Zuber et al., 2018). By consequence, the inward and outward barrier function of the cuticle is severely compromised. In contrast, the envelope structure appears to be normal in *torr* mutant larvae. Nevertheless, the inward and outward cuticle impermeability is also lost in these animals. Assuming that the bidirectional barrier function of the cuticle relies mainly on the integrity of the envelope, we speculate that Torr defines envelope quality without affecting envelope structure. Moreover, these findings indicate that Snu function in envelope construction is normal when Torr function is missing. Hence, Snu and Torr act in parallel in establishing the envelope-based barrier function of the *D. melanogaster* cuticle.

### Torr is needed for CHC deposition on the surface of the cuticle

Previously, we had shown that Snu activity is needed for correct localisation of the extracellular protein Snsl to the pore canals. Snsl, in turn, is needed for the apical distribution of envelope material that displays auto-florescence upon excitation with 405nm light. As pointed out above, these processes are unaffected in *torr* mutant larvae. Thus, Torr is not needed for Snsl trafficking and envelope formation. By contrast, CHC amounts are significantly reduced in wings with reduced *torr* expression. Additionally, we found that Torr-GFP localises to membrane protrusions within the cuticle, reckoning that these structures represent pore canals. We therefore propose that the transporter activity of Torr is required for the amounts of CHC externalized and deposited on the surface of the cuticle via the pore canals. Whether Torr directly transports CHC to the cuticle surface or whether its function is needed indirectly for this process by establishing a structure that facilitates CHC transport remains to be investigated.

### Analogies between vertebrate and invertebrate skin barrier formation

The establishment of a lipid-based ECM in the vertebrate skin involves the function of the ABC transporter ABCA12 (Kelsell, Norgett et al., 2005, Li, Frank et al., 2011). In humans, ABCA12 is involved in the deposition of lipids into the intercellular space during differentiation of the lamellar granules in the skin (Scott, Rajpopat et al., 2013). Mutations in the ABCA12 coding gene cause Harlequin-type ichthyosis (HI), lamellar ichthyosis type 2 (LI) or congenital ichthyosiform erythroderma (CIE), autosomal recessive congenital disorders associated with skin lesions that are lethal shortly after birth (Akiyama, 2014, Akiyama, 2017). Originally, ABCH transporters possibly derived from ABCA transporters during evolution (Dermauw & Van Leeuwen, 2014), and therefore may be lipid transporters, as well. Intriguingly, larvae with eliminated Torr function die shortly after hatching, and CHC levels are reduced in *torr* deficient wings, both paralleling humans with dysfunctional ABCA12. To some extent, hence, the molecular mechanisms of lipid deposition into the skin by an ABC transporter seem to be conserved between vertebrates and invertebrates.

## Materials and Methods

### Fly husbandry and genetics

Flies were cultivated in vials with standard cornmeal-based food at 18 or 25°C. To collect embryos and larvae, flies were kept in cages on apple-juice plates garnished with fresh baker’s yeast at 25°C. Mutations were maintained over balancer chromosomes carrying insertions of GFP expressing marker genes (Dfd-YFP or Kr-GFP). This allowed us to identify homozygous non-GFP embryos or larvae as mutants under a fluorescence stereo-microscope (Leica).

To generate genetically stable mutations in *torr*, the Minos elements in the coding sequence of *torr*^MB00606^ were excised using heat-shock-induced *Minos*-transposase. Larvae and pupae were exposed to daily heat-shocks at 37°C for 1 hour, thereby activating the *Minos*-transposase in the male germ line of the developing flies. Presence or absence of the *Minos* element was determined by the EGFP enhancer trap contained within the 7.5 kb element, which gives a fluorescent signal in the eyes of the flies. 19 Δ*Mi00606* stocks, all representing individual excision events, were established. The stocks were screened for viability of the homozygous Δ*Minos* allele. The primers used to amplify genomic DNA for sequencing of the excision footprint were CCATAGCACGCTCCAAATCA and TGCGCATCATCCACAAGAAG.

### RNAi experiments

To down-regulate *torr* (*CG33970*) expression, the KK-line carrying the hairpin RNA construct with the ID 109988 or the GD-lines carrying the IDs 297 and 7619 constructs, respectively, were crossed to flies harbouring either *69B*-Gal4 (epidermis) or both *da*-Gal4 and *7063*-Gal4 (ubiquitous and maternal Gal4, respectively). To down-regulate *torr* in wings, the KK-line was crossed to flies carrying *nub*-Gal4. As a control, the hairpin RNA construct directed against the midgut *chitin synthase 2* (*CS-2*) with the ID GD 10588 was used.

### Construction of GFP-tagged variants of Torr

Total mRNA was prepared from OregonR wild-type stage 17 embryos (RNeasy Micro Kit, Qiagen) and used to produce cDNA (Superscript III First-Strand Synthesis System, Invitrogen). For amplification of both annotated isoforms of *torr* from this cDNA pool, two alternative 5’ primers, ATGGACGCCGCTGCC and ATGCTGGCAGAGGAATCGC, as well as a single 3’ primer, GCTGCTCAAGTTCAAGAAGGGATAA were used. The products were directly ligated into the pCR8/GW/TOPO vector. Sequencing of the purified vector DNA (Miniprep kit, Qiagen) confirmed the presence of *torr* isoform A and B. Next, LR recombination was used to clone the cDNA into Gateway vector pTGW, which contains a UAS promoter and a *GFP* tag 5’ to the insert. The vector containing *torr* cDNA isoform B failed to amplify in bacteria and was rejected. The vector with the A isoform was amplified in DH5α *E. coli* bacteria, purified (PerfectPrep Endofree Maxi Kit, 5Prime) and sent to Fly-Facility (Clermont-Ferrand, France) for transformation of *D. melanogaster* embryos.

### Quantitative RT-PCR

Total mRNA was isolated from OregonR or Dijon2000 wild-type stage 17 embryos (RNeasy Micro Kit, Qiagen), and total cDNA was prepared (Superscript III First-Strand Synthesis System, Invitrogen). For each embryo collection, at least two independent RNA extractions were assayed three times in parallel. RT-PCR was performed with an iCycler and iTaq SYBR Green Supermix, Biorad. The primers used were GCAATATGTGACCGACGATG and GCGGTACAGCAACTGTGAGA, which amplify a 208 bp fragment common to all *torr* isoforms. The primers TGAGTGACAAAACAGGGATCTT and CGATTCCTCTGCCAGCATTT were used to amplify the short *torr* isoform. As a reference, the primers CGTCGAGGCGGTGTGAAGC and TTAACCGCCAAATCCGTAGAGG that amplify a 195 bp fragment of *histone H4* were used. REST software was used to determine the crossing point differences (DCP value) of individual transcripts in treated (sample) and non-treated (control) embryos (Pfaffl, Horgan et al., 2002). The efficiency (*E*) corrected relative expression ratio of the target gene was calculated using the DCP value of *histone H4* expression according to the equation

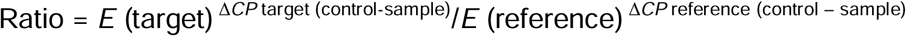

A two-fold change in gene expression was defined as the cut-off for acceptance, as changes up to two-fold are observed also when different wild-type samples are compared (Gangishetti, Breitenbach et al., 2009).

### Microscopy

For transmission electron microscopy (TEM), embryos and larvae were treated and analysed on a Philips CM-10 electron microscope as described in detail previously (Moussian & Schwarz, 2010). Bright field, differential interference contrast (DIC) and fluorescence microscopy were performed with a Nikon eclipse E1000 or a Zeiss Axioplan 2 microscope. Confocal imaging was performed with Zeiss LSM 710 Axio Observer, Bio-Rad Radiance 2000 or Leica TCS SP2. Figures were prepared using the Adobe Photoshop CS6 and Adobe Illustrator CS6 software.

### Chemical analysis of CHCs

To extract cuticular hydrocarbons, 6-hour or 5-day old flies were frozen for 5 min at − 20°C just before removing the wings using micro-scissors. Each pair of wings was immersed for 10 min at room temperature into vials containing 20 μL of hexane. The solution also contained 3.33 ng/µl of C26 (*n*-hexacosane) and 3.33 ng/µl of C30 (*n*-triacontane) as internal standards. After removing the wings, the extracts were stored at −20°C until analysis. All extracts were analyzed using a Varian CP3380 gas chromatograph fitted with a flame ionization detector, a CP Sil 5CB column (25 m × 0.25 mm internal diameter; 0.1 μm film thickness; Agilent), and a split–splitless injector (60 ml/min split-flow; valve opening 30 s after injection) with helium as carrier gas (velocity = 50 cm/s at 120 °C). The temperature program began at 120°C, ramping at 10°C/min to 140°C, then ramping at two °C/min to 290°C, and holding for 10 min. The chemical identity of the cuticular hydrocarbons was checked using gas chromatography-mass spectrometry system equipped with a CP Sil 5CB column (Everaerts et al., 2010). The amount (ng/insect) of each component was calculated based on the readings obtained from the internal standards. For the sake of clarity we only show the principal CH groups: the overall CHs sum (∑CHs), the sum of desaturated CHs (∑Desat), the sum of linear saturated CHs (∑Lin) and the sum of branched CHs (∑Branched).

## Acknowledgments

This work was supported by the German Research Foundation (DFG, MO1714/9-1) and the National Science Foundation of China (NSFC, 31761133021).

## Figure legends

*Figure S1*

Represented by protein sequences from the hemimetabolous insect *Locusta migratoria* and the holometabolous insect *D. melanogaster*, insect species have three H-type ABC transporters. The ancestral protein that is shared with crustaceans (*Daphnia*), is ABCH-9A (CG11147 in *D. melanogaster*).

*Figure S2*

Torr-GFP expressed in the epidermis (*69B*-Gal4, UAS-*torrGFP*) or ubiquitously (tub-Gal4, UAS-*torrGFP*) normalises the cuticle impermeability in *torr*^Δ*MiM18*^ homozygous mutant larvae (compare to Fig. 3).

*Figure S3*

Wings of control flies expressing hpRNA against midgut chitin synthase 2 transcripts (*nub*-Gal4 × UAS-*CS-2*^*wgIR*^, *CS-2*^*wgIR*^) are impermeable to Eosin Y until 50°C. At 55°C, the dye penetrates the posterior margin of the wing of these flies. Penetration is more pronounced at higher temperatures (60°C). Wings of flies expressing hpRNA against *torr* (*nub*-Gal4 × UAS-*KK109988, torr*^*wgIR*^) transcripts are impermeable to Eosin Y until 45°C. Eosin Y penetrates the posterior half of the wing of these flies at 50°C, and the whole wing at 55°C.

