## Supplementary figures and images for "Dysfunction of Torr causes a Harlequin-type ichthyosis-like phenotype in *Drosophila melanogaster*"

### Supplementary figure 1

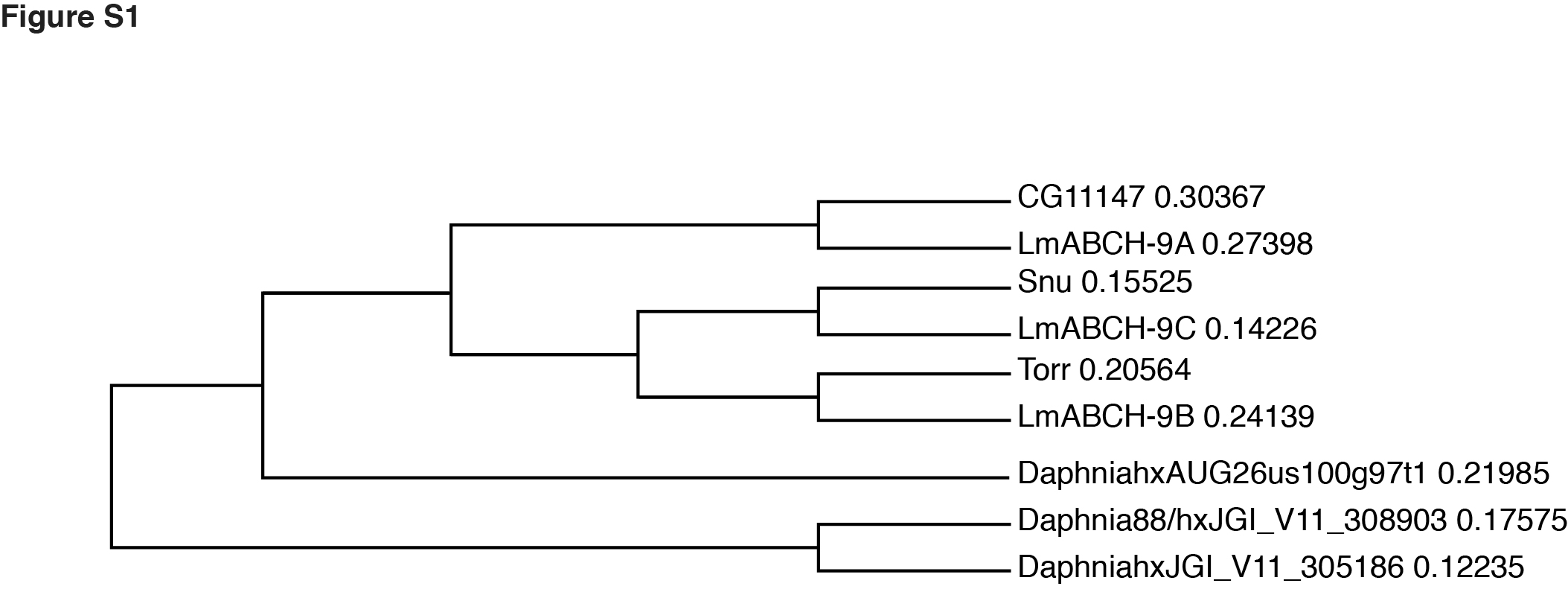

### Supplementary figure 2

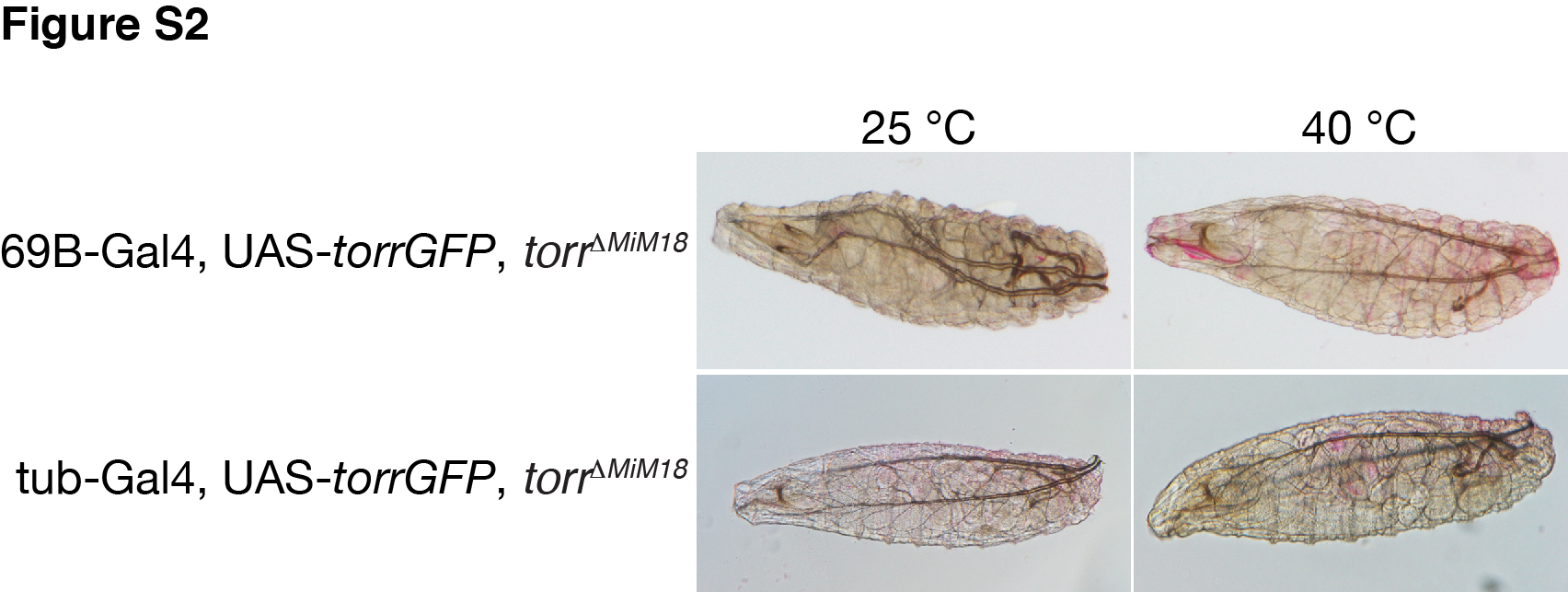

### Supplementary figure 3

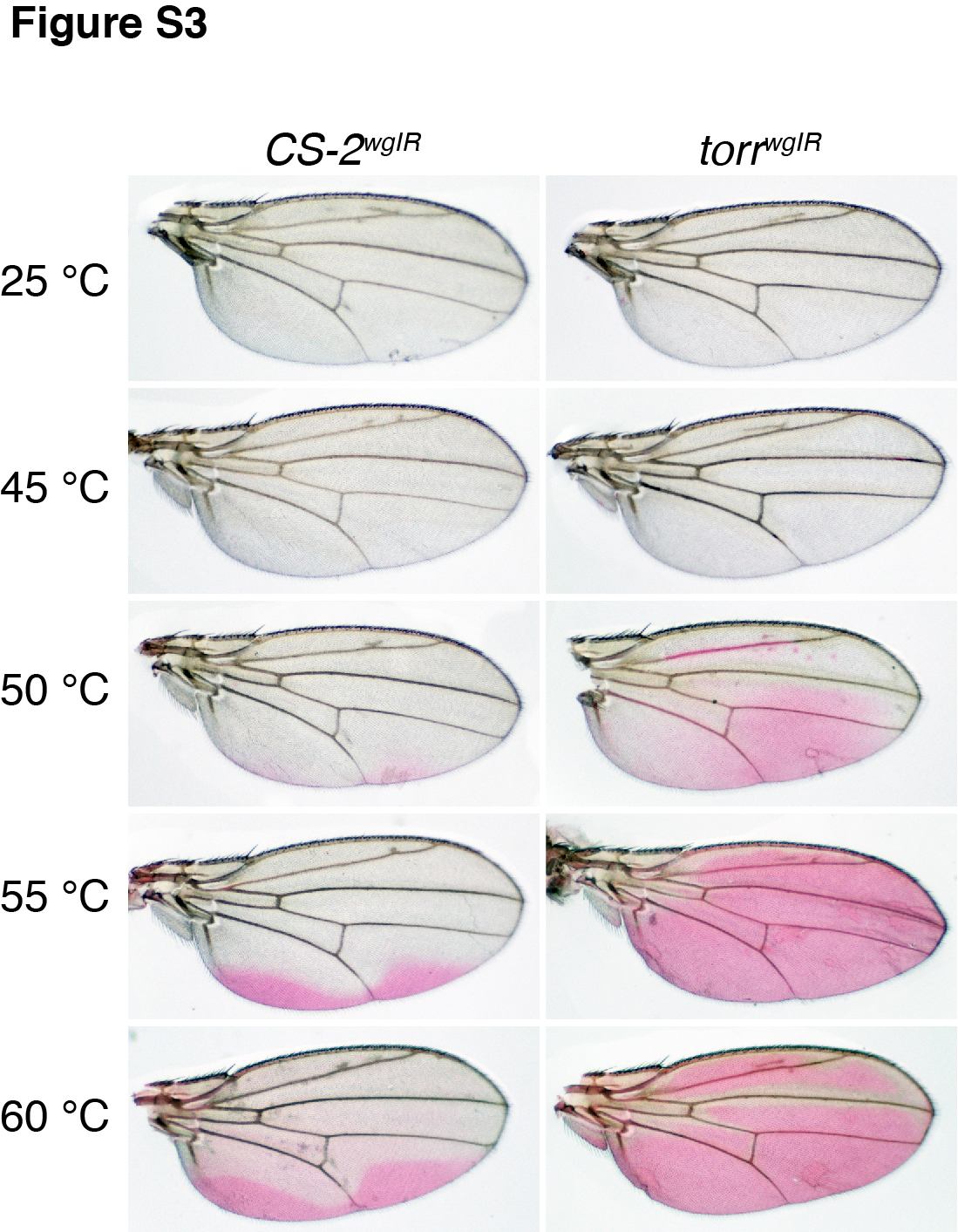
